# The Contribution of Audition and Proprioception in Unisensory and Multisensory Target Reaching

**DOI:** 10.1101/2025.07.02.662728

**Authors:** Ivan Camponogara

**Author notes:** **Corresponding author:** Ivan Camponogara, Haptic, Auditory, Kinematic Laboratory (HAKi Lab), Department of Psychology, College of Natural and Health Sciences, Zayed University Abu Dhabi, United Arab Emirates. All data and data analysis scripts are available at the following link: https://osf.io/cu2q3/.

## Abstract

Everyday actions often involve reaching for targets sensed by auditory and proprioceptive senses (reaching a ringing smartphone in the dark or tapping on it while holding it with the other hand). However, it is still unclear whether reaching performance toward auditory and proprioceptive targets is modality-specific and whether performance improves under multisensory compared to unisensory conditions. Here, we addressed these questions by measuring reaching performance toward auditory, proprioceptive, and combined audio-proprioceptive targets. Accuracy was generally similar across conditions, but precision was lower for auditory targets compared to proprioceptive and audio-proprioceptive targets. A second experiment investigated whether providing additional proprioceptive information while reaching for an auditory target could improve precision in subsequent auditory-only trials. A slight improvement was observed, indicating that proprioceptive cues may help reduce spatial variability in auditory-guided actions, though not to the level seen in the multisensory condition. Overall, the results suggest that while the target modality slightly impacts movement accuracy, it has a significant impact on movement precision, with proprioceptive input playing a crucial role in enhancing precision. The concurrent availability of auditory and proprioceptive target information does not enhance precision beyond that achieved with proprioceptive information alone, whereas proprioception can modestly improve subsequent auditory-guided reaching.

## Introduction

In everyday life, humans constantly use auditory and proprioceptive information to localize objects and execute goal-directed actions. For instance, when silencing a smartphone alarm upon waking, we use auditory cues to locate and reach the smartphone in the dark and rely on proprioceptive feedback from the hand holding it or on both modalities (i.e., proprioception and the ringing sound) to tap on it with the other hand and silence it. Although both sensory modalities support the same goal (i.e., successfully reaching a target), the mechanisms underlying their sensorimotor processes remain poorly understood. According to the perceptual mechanisms characterizing each sensory modality, the sensorimotor processes involved in auditory and proprioceptive target reaching may be modality-specific. Auditory localization primarily relies on interaural time and level differences, which enable relatively accurate identification of sound sources along the azimuth (i.e., horizontal) dimension (Middlebrooks, 2015; Middlebrooks and Green, 1991; Makous and Middlebrooks, 1990; Wightman and Kistler, 1990; Kato et al, 2003). However, localization accuracy diminishes along the depth dimension, where sound distance is often misjudged (Middlebrooks, 2015). Proprioceptive localization, such as identifying the position of an unseen limb, depends on spatial encoding in joint-centered coordinates (van Beers et al, 1998). Proprioceptive judgments tend to be biased in both azimuth and depth, though exhibiting higher precision in the depth dimension than in azimuth (Kuling et al, 2016; Liu et al, 2018; van Beers et al, 1998, 1996, 1999a, 2002b, 1999b, 2002a; Camponogara and Volcic, 2021). Taken together, these studies suggest that actions toward auditory and proprioceptive targets may show modality-specific patterns of accuracy and precision, shaped by the distinct encoding strategies of each sensory system.

However, several inconsistencies arise when comparing findings across studies that independently investigate reaching movements within different sensory modalities. Some studies on proprioceptive target reaching suggest more accurate and precise movements (Jones et al, 2012; Monaco et al, 2010; McGuire and Sabes, 2009; Blouin et al, 2014; Cameron and Ĺopez-Moliner, 2015) compared to those toward auditory targets (Parseihian et al, 2014; Maće et al, 2012; Boyer et al, 2013), across both azimuth and depth dimensions. In contrast, other studies suggest shared accuracy or precision along either the azimuth or depth dimension (Mikula et al, 2018; Flannigan et al, 2018; Khanafer and Cressman, 2014). This points to a potentially overlapping mechanism underlying target-reaching across sensory modalities. Nonetheless, such conclusions should be interpreted with caution, as these studies have not directly compared actions toward auditory and proprioceptive targets within the same experimental context. The limited number of studies comparing reaching movements toward auditory and proprioceptive targets suggests that proprioceptive-guided actions tend to be more accurate and precise than auditory-guided actions (Cuppone et al, 2018, 2019). Notably, in these studies, proprioceptive target reaching involved relocating a limb to a previously sensed position. Participants were first guided, either actively or passively, from a start location to a target, after which the limb was withdrawn and subsequently moved to reproduce the remembered location. This task relies not only on proprioceptive input but also on sensorimotor memory of the preceding movement, (Soechting and Flanders, 1989b,a; Tillery et al, 1991; Darling and Miller, 1993; Berkinblit et al, 1995; Pettypiece et al, 2009; Jones and Henriques, 2010) which has been shown to significantly enhance both accuracy and precision (Jones et al, 2012). Integrating proprioceptive feedback with sensorimotor memory likely enhances spatial estimation, potentially explaining the observed advantage of proprioceptive over auditory-guided reaching in these paradigms. Furthermore, in these studies, movement was often constrained on a rail along one movement dimension (i.e., oblique, 45 degrees), which likely reduced the natural variability inherent in skilled motor behavior (Latash, 2012). Thus, discrepancies among studies, combined with the limited direct experimental comparisons between actions toward auditory and proprioceptive targets, limit our ability to conclude whether reaching movements are modality-specific.

A further open question concerns whether a multisensory advantage occurs when both sensory modalities are simultaneously available. Studies in the visuo-proprioceptive domain showed improved action performance in the multisensory condition compared to each of the unisensory conditions (visual and proprioceptive) (Camponogara, 2023). However, in the audio-proprioceptive domain, it remains unclear whether the concurrent availability of both auditory and proprioceptive target information enhances motor performance. Previous studies addressing multisensory-motor training on auditory localization showed that the multisensory-motor association between proprioceptive and motor signals from the reaching arm and the auditory target improves subsequent auditory localization (Cuppone et al, 2018, 2019; Valzolgher et al, 2020a,b; Lombera et al, 2022; Valzolgher et al, 2023a,b, 2024). This suggests that the concurrent availability of auditory and proprioceptive inputs may lead to better performance than each unisensory input alone. In contrast, studies on auditory-proprioceptive localization suggest a strong influence of arm proprioception on auditory perception when localizing a handheld speaker (Lackner and Shenker, 1985). It is thus plausible to think that, in multisensory contexts, actions are primarily guided by proprioceptive information.

To address these questions, we conducted an experiment in which participants performed reaching movements toward targets defined by audition, proprioception, or both sensory modalities, presented at three azimuthal locations. Performance was evaluated in terms of both accuracy and precision along the azimuth and depth dimensions to assess the relative contribution of each sensory modality to action guidance and to determine whether their concurrent availability confers a performance benefit. Based on prior findings, two hypotheses can be advanced: 1) Actions toward auditory and proprioceptive targets may exhibit similar accuracy and precision along specific movement dimensions (van Beers, 2009; Parseihian et al, 2014; Maće et al, 2012; Boyer et al, 2013; Flannigan et al, 2018; Khanafer and Cressman, 2014; van Beers et al, 1996, 1999a, 2002b, 1999b, 2002a). In contrast, 2) actions toward proprioceptive targets may be more accurate and precise across both dimensions (Jones et al, 2012; Monaco et al, 2010; McGuire and Sabes, 2009; Blouin et al, 2014; Cameron and Ĺopez-Moliner, 2015). Furthermore, if a multisensory advantage exists (Cuppone et al, 2018, 2019; Valzolgher et al, 2020a,b; Lombera et al, 2022; Valzolgher et al, 2023a,b, 2024), we expected that reaching in the combined audio-proprioceptive condition would exhibit higher accuracy and precision than either unisensory condition.

Conversely, it may be that, in the multisensory condition, motor performance may rely on proprioceptive target information (Lackner and Shenker, 1985), thus leading to similar action performance as in proprioceptive target reaching.

## Experiment 1

### Material and Method

#### Participants

We tested twenty-five participants (right-handed, 1 male, age 21.3 *±* 1.9 years). All self-reported normal hearing and no known history of neurological disorders. All of the participants were näıve to the purpose of the experiment. The experiment was undertaken with the understanding and informed written consent of each participant, and the experimental procedures were approved by the Institutional Review Board of Zayed University, Abu Dhabi, in compliance with the Code of Ethical Principles for Medical Research Involving Human Subjects of the World Medical Association (Declaration of Helsinki).

#### Apparatus

The custom-made experimental apparatus consisted of a perforated wooden board of 600 by 600 mm supported by four 100 mm high pedestals placed on the experimental table. Three speakers (Gikfunk, 4O 0hm 3 Watt, 40 mm diameter, 30 mm height) were placed underneath the board, powered through a USB cable, and connected with an external soundcard (Motu 4Pre) via a mini amplifier (PAM8403). The center of the speakers was placed 250 mm far at 0 (i.e., in front), 45, and 90 degrees on the left with respect to a 5 mm high rubber bump that acted as a starting point. The bump was placed 100 mm far from the side of the wooden board closest to the subject position (Figure 1 A). A 5 mm high rubber bump was placed underneath the center of each speaker and was used as a reference for the left index position during the proprioceptive and audio-proprioceptive conditions. The right-hand index finger movement and the speakers’ positions were acquired at 60 Hz with a resolution of 720 × 1280 pixels using a webcam (Logitech BRIO Ultra) placed 600 mm over the wooden board with a camera tripod. The x and y pixel coordinates of the finger’s movement and speakers’ positions were extracted off-line by using DeepLabCut (Nath et al, 2019), a pose estimation toolbox system in Python. Finally, a chin rest was placed on the front side of the wooden board to avoid head movements during the experiment. The chin position was set at 25 cm height from the wooden surface, and the chair height was adjusted at the participants’ comfortable height. A pink noise of 1000 ms duration (65 db) was used as the auditory target, whereas a 100 ms duration pure tone of 500 Hz (65 dB) was used to signal the end of the trial. The same tone was used to signal the start of the trial in the proprioceptive condition. Pure tones were delivered through two external speakers (Logitech 980-000816 Z150). The camera and sound delivery were controlled via MATLAB (MathWorks), which was installed on a Dell Alienware laptop.

**Figure 1:**
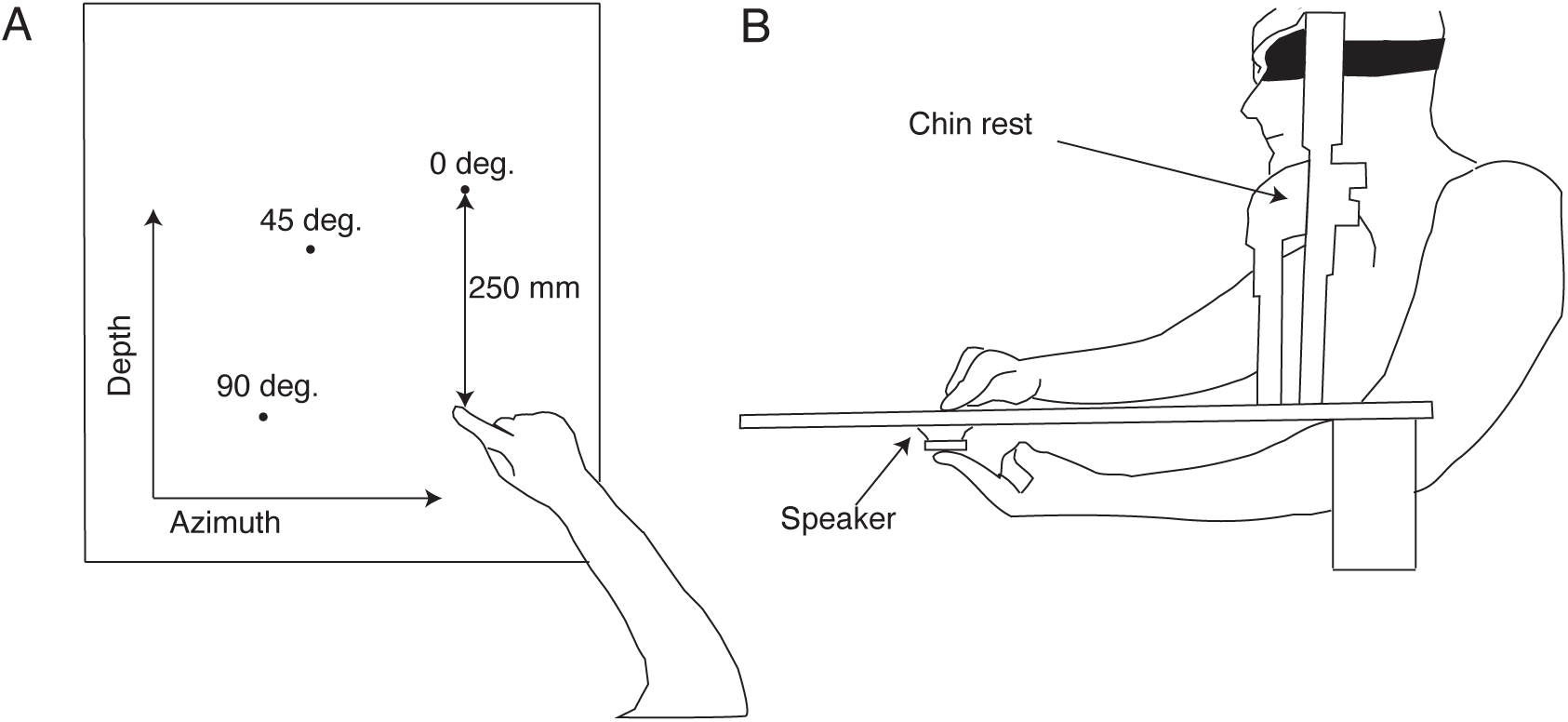
Experiment setup. A) Top view: Targets were positioned at 250 mm in front (0 degrees) or laterally (45 and 90 degrees) from a start position. B) Side view in P and AP condition: Participants placed the left index underneath the center of one of the three speakers. As soon as they heard the start sound (P) or the pink noise (AP), participants were instructed to reach the tip of their left finger (P condition) or the point where they heard the sound and felt the finger position (AP condition).

#### Procedure

Participants were blindfolded for the whole duration of the experiment. All the trials started with the participant’s head on the chin rest, the index digit of the right hand positioned on the start position, and the left hand on the left side of the chin rest. In an Auditory condition (A), the pink noise was delivered, and participants had to reach the point where they felt the noise coming from. In a Proprioceptive (P) and Audio-Proprioceptive (AP) condition, before each trial, participants were instructed to move the index of the left hand underneath the specified target speaker over the bump placed at the center of it. After 1 second, in the P condition, the start sound was delivered from the external speakers, whereas in AP, the pink noise was delivered from the speaker over the left finger. As soon as participants heard the sound (P) or the pink noise (AP), they had to reach the point where they thought the tip of their left-index finger was or both the tip of the finger and the pink noise were (Figure 1 B). In all the sensory conditions, after 2 seconds from the pink noise or the start tone, the end tone was delivered, and participants had to move their right hand back to the start position and their left hand back to the side of the chin rest in P and AP conditions. Before each experiment, participants underwent a practice phase in all the conditions, consisting of 10 trials, where they got accustomed to the task. We performed 20 trials for each target, leading to 60 trials per condition (180 trials per participant). Conditions were pseudo-randomized, with A always being the first. Then, half of the participants underwent the P and AP order, and the other half the AP and P order. This randomization was chosen to prevent trials in A condition to be affected by any multisensory learning effect if AP and P conditions were performed at first (Cuppone et al, 2018, 2019; Valzolgher et al, 2020a,b, 2023b,a, 2024).

#### Data analysis

Kinematic data were analyzed in R R Core Team (2020). The x and y coordinates of the index and speakers’ position in pixels were smoothed and differentiated with a third-order Savitzky-Golay filter with a window size of 21 points. These filtered data were then used to compute velocities and accelerations of the Index finger in two-dimensional space. Movement onset was defined as the moment of the lowest, non-repeating Index finger acceleration value prior to the continuously increasing Index finger acceleration values, while the end of the movement was defined by applying the same algorithm but starting from the end of the recorded values Camponogara and Volcic (2021). Subject 7 voluntarily withdrew from the experiment when in the P condition, performing 32 trials in total instead of 60. From the 4348 trials, we discarded from further analysis the trials in which the camera (48 trials) or the algorithm (137 trials) did not capture the end of the movement. The exclusion of these trials (185 trials, 4.2% of all trials) left us with 4163 trials for the final analysis. For each trial, we calculated the endpoint positions along the azimuth and depth dimensions (i.e, the x and y coordinates).

We focused our analyses on the x and y coordinates of the endpoint error, defined as the distance between the center of the target and the index’s landing point. The x and y endpoint error coordinates were analyzed using the Bayesian multivariate linear mixed-effects distributional regression model, estimated using the brms package Bürkner (2017), which implements Bayesian multilevel models in R using the probabilistic programming language Stan Carpenter et al (2017). The distributional regression model allowed us to estimate the posterior distribution of both the accuracy and the SD, taking into account both within and between-subject variability Camponogara and Volcic (2021). The model included as predictors (i.e., fixed effects) the categorical variable Condition (A, P, and AP) and the continuous variable Direction (0, 45, and 90 degrees). Thus, the model estimated the average accuracy in depth and azimuth ( *β_Condition_*), its change as a function of the target direction (*β_Direction_*, slope), the average SD in depth and azimuth *σ_Condition_*, and the SD change in azimuth and depth as a function of the target direction (*σ_Direction_*, slope). The model included the independent random (group-level) effects for subjects.

The model was fitted considering weakly informative prior distributions for each parameter to provide information about their plausible scale. We used Gaussian priors for the Condition fixed-effect predictor (*β_Condition_*: mean = 0 and sd = 80, *β_Direction_*: mean = 0 and sd = 5, *σ_Condition_*: mean = 2.5 and sd = 2,*σ_Direction_*: mean = 0 and sd = 2), whereas for the group-level standard deviation parameters we used zero-centered Student *t*-distribution priors, with scale values defined by the “get prior” function. (azimuth = *df* = 3, scale = 32.9, depth = *df* = 3, scale = 34.5). Finally, we set a prior over the correlation and residual correlation matrix that assumes that smaller correlations are slightly more likely than larger ones (LKJ prior set to 2).

We ran four Markov chains simultaneously, each for 4,000 iterations (1,000 warm-up samples to tune the MCMC sampler) with the delta parameter set to 0.99 for a total of 12,000 post-warm-up samples. Chain convergence was assessed using the *R*^^^ statistic (all values equal to 1) and visual inspection of the chain traces. Additionally, the predictive precision of the fitted models was estimated with leave-one-out cross-validation by using the Pareto Smoothed Importance Sampling (PSIS). We found 1 pareto k value higher than 0.7, and re-fit the model using the reloo function in brms Bürkner (2017). After the model re-fit, all the pareto k values were lower than 0.7.

The posterior distributions we have obtained represent the probabilities of the parameters conditional on the priors, model, and data, and they represent our belief that the “true” parameter lies within some interval with a given probability. We summarize these posterior distributions by computing the medians and the 95% Highest Density Intervals (HDI). The 95% HDI specifies the interval that includes, with a 95% probability, the true value of a specific parameter. To evaluate the differences between the two compared conditions, we have simply subtracted the posterior distributions of *β_Condition_*, *β_Direction_*, *σ_Condition_*, and *σ_Direction_* between specific conditions. The resulting distributions are denoted as the credible difference distributions and are again summarized by computing the medians and the 95% HDIs.

For statistical inferences about the *β_Condition_*, *β_Direction_*, *σ_Condition_* and *σ_Direction_* we assessed the overlap of the 95% HDI with zero. A 95% HDI that does not span zero indicates that the accuracy (*β_Condition_*) and SD (*σ_Condition_*) in depth and azimuth were credibly different than zero in that specific condition. Additionally, a 95% HDI that does not span the zero for the *β_Direction_* and *σ_Direction_* indicates that both the accuracy and SD, respectively, are modulated according to the target direction. For statistical inferences about the differences between conditions, we applied an analogous approach. A 95% HDI of the credible difference distribution that does not span zero is taken as evidence that the accuracy (*β_Condition_*) or SD (*σ_Condition_*) in the two conditions differ from each other. The same applied to the predictor Direction. To assess the strength of the evidence, we computed the Bayes factor for each comparison. The reported Bayes factor values (BF_10_) that are higher than 3 provide evidence in support of a difference between conditions, whereas values below ^1^ provide evidence in support of an absence of a difference between conditions Lee and Wagenmakers (2014).

## Results

In general, we found an overall target undershoot across conditions (Figure 2A, Figure 3A). The accuracy was credibly similar between conditions (Figure 3B), with a minimal difference between A-AP (depth = −27 pixels, HDI = −49.7, −4.67 pixels; azimuth = −3 pixels, HDI = −18.8, 12.2 pixels) and A - P (depth = −27 pixels, HDI = −51.3, −4.47 pixels; azimuth = −5 pixels, HDI = −20.2, 9.41 pixels) along the depth dimension. However, all BF_10_ were *<*3, indicating anecdotal evidence toward the alternative hypothesis. The slope analysis revealed a credible similar accuracy across target directions for P and AP (i.e., flat slope), whereas was slight increase in the accuracy along the depth dimension for the A condition, with, however, an anecdotal evidence toward the alternative hypothesis (BF_10_ = 0.3). No credible difference for the slope was found between conditions.

**Figure 2:**
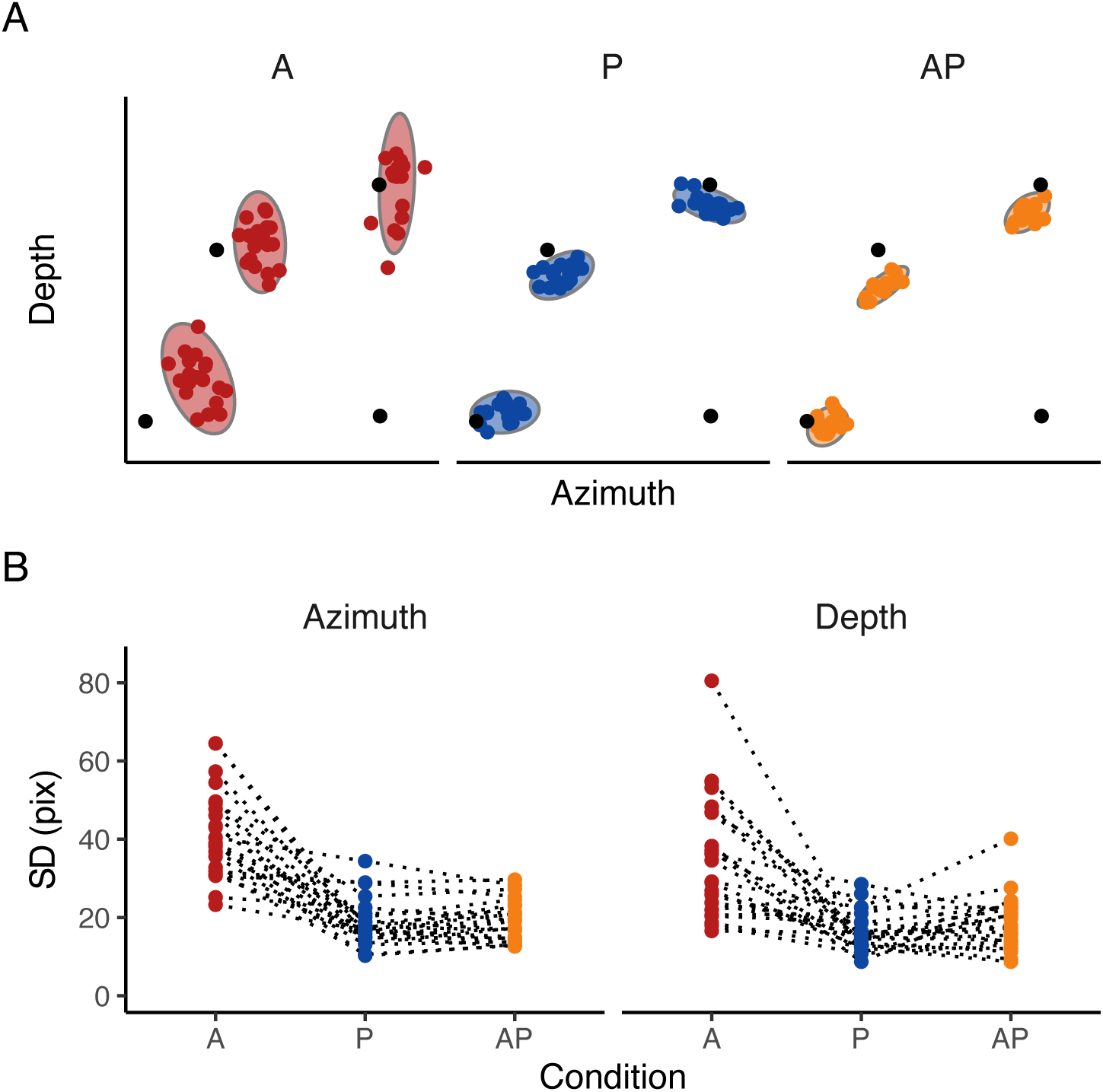
Raw data. A) Final landing points (dots) and their 95% confidence ellipses standard deviation of a representative participant for experiment 1. The bottom right black dot represents the positions of the starting point, whereas the other three dots represent the center of the three targets. B) Raw SD. Raw SD of each participant in A, P, and AP conditions in the azimuth and depth dimensions. Dotted lines connect each participant’s result in each condition.

**Figure 3:**
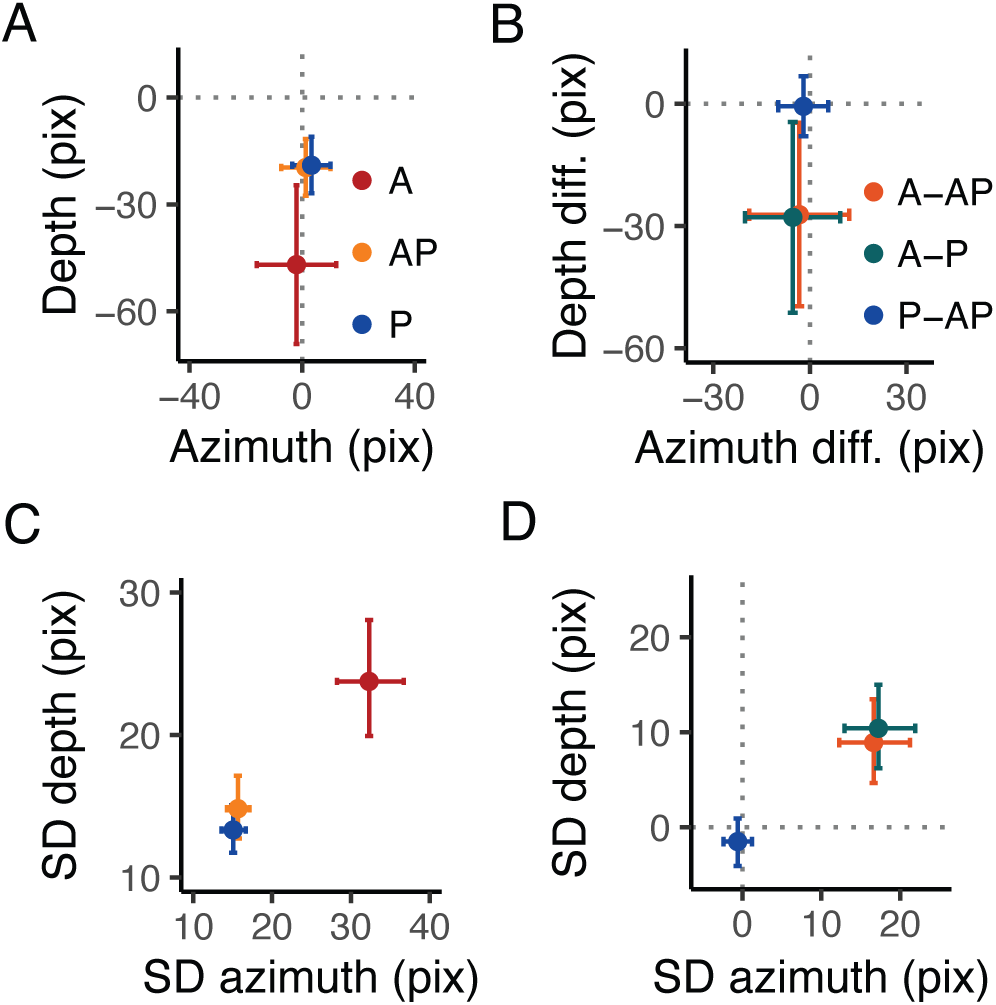
Experimental results. A) Estimates of the accuracy in the azimuth and depth. The dots represent the mean, and the error bars denote the 95% HDI of the estimates. The intersection of the grey dashed lines represents the center of the targets. In A, positive values in the depth dimension indicate an overshoot, whereas positive values in the azimuth indicate that the index finger landed on the right side of the target. B) Estimates of the differences in accuracy in the azimuth and depth between conditions. The dots represent the mean, and the error bars denote the 95% HDI of the estimates. The grey dashed line represents the points of equal error in the depth and azimuth dimensions. C) Estimates of the SD in the azimuth and depth. The dots represent the mean, and the error bars denote the 95% HDI of the estimates. The intersection of the grey dashed lines represents the center of the targets. D) Estimates of the differences in SD in the azimuth and depth between conditions. The dots represent the mean, and the error bars denote the 95% HDI of the estimates. The grey dashed line represents the points of equal variability in the depth and azimuth dimensions.

The SD (Figure 2B, Figure 3C and D), instead, was credibly lower with strong evidence toward the alternative hypothesis in A compared to AP in both the depth (BF_10_ *>*100) and azimuth directions (BF_10_ *>*100). The same held for the comparisons between the A and P conditions in both depth and azimuth (BF_10_ *>*100). The comparisons between P and AP revealed a credibly similar SD (all BF_10_ *<*1). The slope showed a minimal effect of target direction on SD, showing a slight decrease of SD in A and a slight increase in P as the target is laterally displaced (i.e., 90 degrees) along the depth dimension, whereas along the azimuth in P and AP. However such changes were irrelevant (all BF_10_ *<*1). The analysis between conditions showed a minimal difference in slope between H and AP and between A and P in the depth (P-AP: 0.04 pixels, HDI = 0.004, 0.08 pixels, A-P: −0.09 pixels, HDI = −0.15, 0.03 pixels) dimension, and between A and AP, and A and P in the azimuth dimension (A-AP: 0.15 pixels, HDI = 0.004, 0.3 pixels, A-P: 0.15 pixels, HDI = 0.01, 0.3 pixels). However, all BF_10_ were *<*1, indicating anecdotal evidence toward the alternative hypothesis.

### Conclusion Experiment 1

Taken together, results showed a credibly similar accuracy in A, P, and AP, confirming a shared accuracy across conditions (Parseihian et al, 2014; Maće et al, 2012; Valzolgher et al, 2025; Boyer et al, 2013; Haggard et al, 2000; Jones et al, 2010; van Beers et al, 1998) in the azimuth dimension, and a slightly higher bias for the depth dimension in auditory target reaching (Middlebrooks, 2015; Zahorik and Anderson, 2015). Actions toward proprioceptive and audio-proprioceptive targets were more precise than those toward auditory targets, hinting at a higher reliance on proprioception for action consistency when reaching multisensory targets. Such a process raises the question of the effectiveness of additional proprioceptive target information in ameliorating subsequent reaching performance toward an auditory target. Several studies in the literature reported a better auditory target localization following sensorimotor (proprioceptive and motor) training, where participants actively or passively reached an auditory target (Cuppone et al, 2018, 2019; Valzolgher et al, 2020a,b, 2023b,a, 2024; Hüg et al, 2019, 2022). Such studies suggested that the improved performance after the training was due to a cross-modal effect, where sensorimotor inputs from the reaching arm are used to map the auditory target location. In other words, the improved auditory localization performance after the auditory target reaching may be due to the combination of proprioceptive and motor inputs signaling the final extension of the reaching arm with the auditory target location (Cuppone et al, 2018; Valzolgher et al, 2020a). It is thus still unclear whether adding proprioceptive target information may ameliorate a subsequent action performance toward an auditory target. To investigate the effect of additional proprioceptive information on auditory target reaching, we performed a second experiment where we measured the auditory target reaching accuracy and precision before (Pre) and after (Post) reaching an audio-proprioceptive target. If participants use proprioception to map the auditory target location, results will show a better action performance (higher accuracy and precision) in the Post compared to the Pre condition. In contrast, a lack of action improvement would prove that additional proprioceptive target information is insufficient to map an auditory target location.

## Experiment 2

### Material and Method

#### Participants

We tested twenty-five participants (right-handed, 1 male, age 25.1 *±* 8.9 years). All had normal or corrected-to-normal vision and no known history of neurological disorders. All of the participants were näıve to the purpose of the experiment. The experiment was undertaken with the understanding and informed written consent of each participant, and the experimental procedures were approved by the Institutional Review Board of Zayed University, Abu Dhabi, in compliance with the Code of Ethical Principles for Medical Research Involving Human Subjects of the World Medical Association (Declaration of Helsinki).

#### Apparatus and Procedure

The apparatus and procedures were the same as Experiment 1. Participants performed an A condition, named Pre, an AP condition, and a second A condition, named Post. We run 20 trials for each target, leading to 60 trials per condition (180 trials per participant).

#### Data analysis

The raw data processing and statistical analysis were the same as in Experiment 1. From the 4500 trials, we discarded from further analysis the trials in which the end of the movement was not captured correctly by the camera (213 trials in total) or the algorithm (73 trials in total). The exclusion of these trials (286 trials, 6.3% of all trials) left us with 4214 trials for the final analysis. For each trial we calculated the endpoint positions along the azimuth and depth dimensions. As in Experiment 1, we focused our analyses on the endpoint error. The *R*^^^ and visual inspection of the chain traces confirmed successful chains convergence. All pareto k values were lower than 0.7.

A further analysis was performed to define whether a learning effect occurred throughout each condition. For each participant, we split the 60 trials in 3 blocks of 20 trials each. We then analyzed the endpoint error using the Bayesian multivariate linear mixed-effects distributional regression model, with the categorical variable Condition (Pre, AP, Post) in combination with the centered continuous variable Block (1,2,3). The model included the independent random (group-level) effects for subjects. The model was fitted considering weakly informative prior distributions for each parameter to provide information about their plausible scale. We used Gaussian priors for the Condition and Block fixed-effect predictors (*β_Condition_*: mean = 0 and sd = 80, *σ_Condition_*: mean = 2.5 and sd = 2, *β_Block_*: mean = 0 and sd = 5, *σ_Block_*: mean = 0 and sd = 2), whereas for the group-level standard deviation parameters we used zero-centered Student *t*-distribution priors, with scale values defined by the “get prior” function. (azimuth = *df* = 3, scale = 43.8, depth = *df* = 3, scale = 41). Finally, we set a prior over the correlation and residual correlation matrix that assumes that smaller correlations are slightly more likely than larger ones (LKJ prior set to 2). The *R*^^^ and visual inspection of the chain traces confirmed successful chains convergence. We found 1 pareto k value higher than 0.7, and re-fit the model using the reloo function in brms (Bürkner, 2017). After the model re-fit, all the pareto k values were lower than 0.7. We focused our analysis on the *σ_Condition_*parameter, which corresponds to the within-participant SD in azimuth and depth dimensions in each Condition, and the estimates of the Block parameter *σ_Block_*, which corresponds to the change of within-participant SD along the azimuth and depth dimension as a function of the Block (i.e., the slope). The reason for focusing on the model’s *σ* component is rooted in the findings of Experiment 1, where it was observed that providing proprioceptive target information resulted in a significant reduction in the SD.

## Results

### Accuracy and SD

In general, we found an overall target undershoot across conditions (Figure 4A, Figure 5A). The accuracy was credibly similar between conditions (all BF_10_ *<*3, Figure 5 B) and did not vary according to the target directions (all BF_10_ *<*2). As for experiment 1, there was a tendency of a higher target undershooting in the Pre compared to the AP condition along the depth dimension (Pre - AP = −19.3 pixels, HDI = −38.9, 0.1 pixels), but it was not credibly different. Even though the slope in the Pre condition was credibly higher than in the AP condition (difference = 0.3, HDI = 0.1, 0.6), we found anecdotal evidence toward the alternative hypothesis (BF_10_ = 0.66).

**Figure 4:**
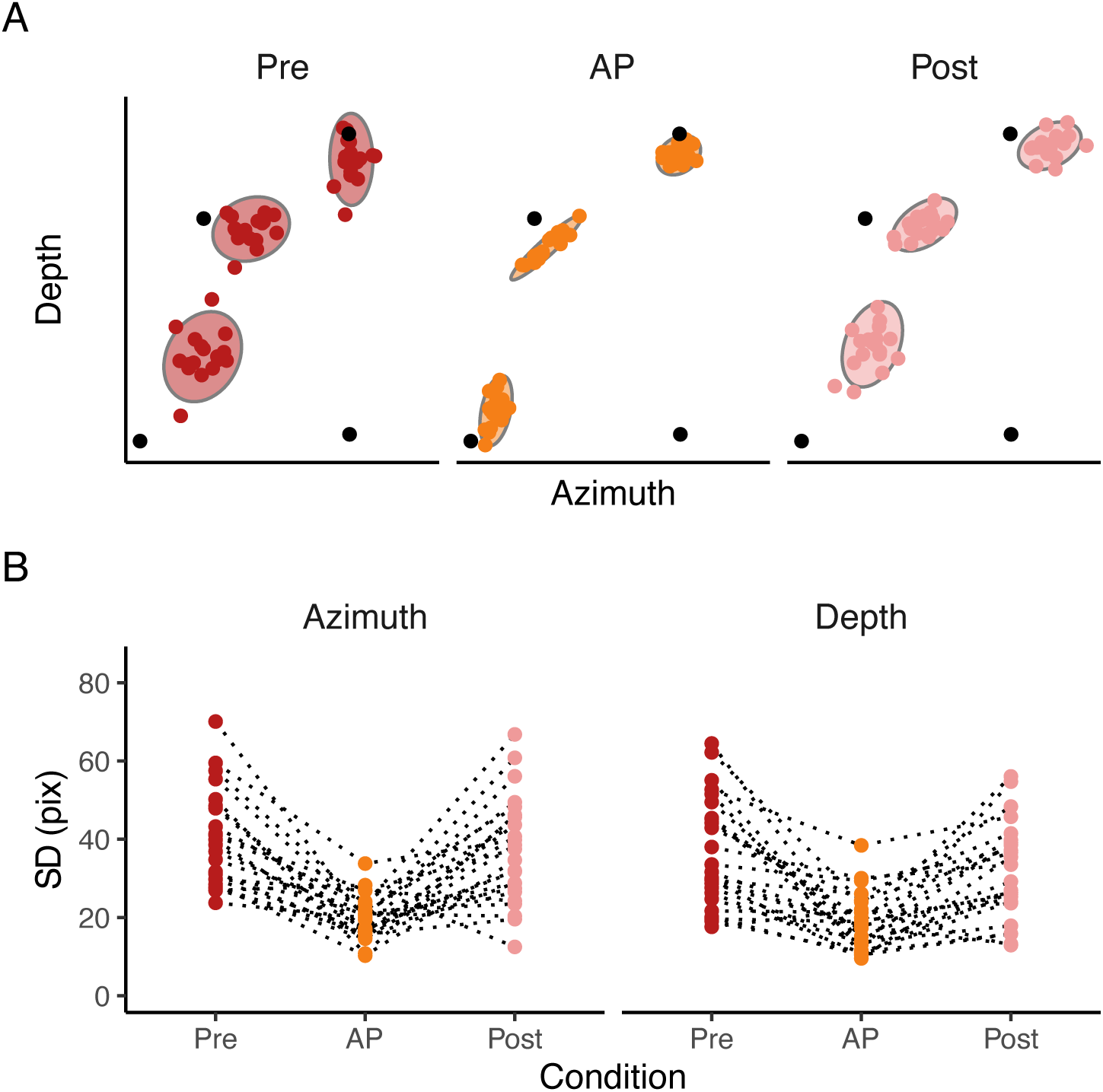
Raw data. A) Final landing points (dots) and their 95% confidence ellipses standard deviation of a representative participant for experiment 2. The bottom right black dot represents the positions of the starting point, whereas the other three dots represent the center of the three targets. B) Raw SD. Raw SD of each participant in Pre, AP, and Post conditions in the azimuth and depth dimensions. Dotted lines connect each participant’s result in each condition.

**Figure 5:**
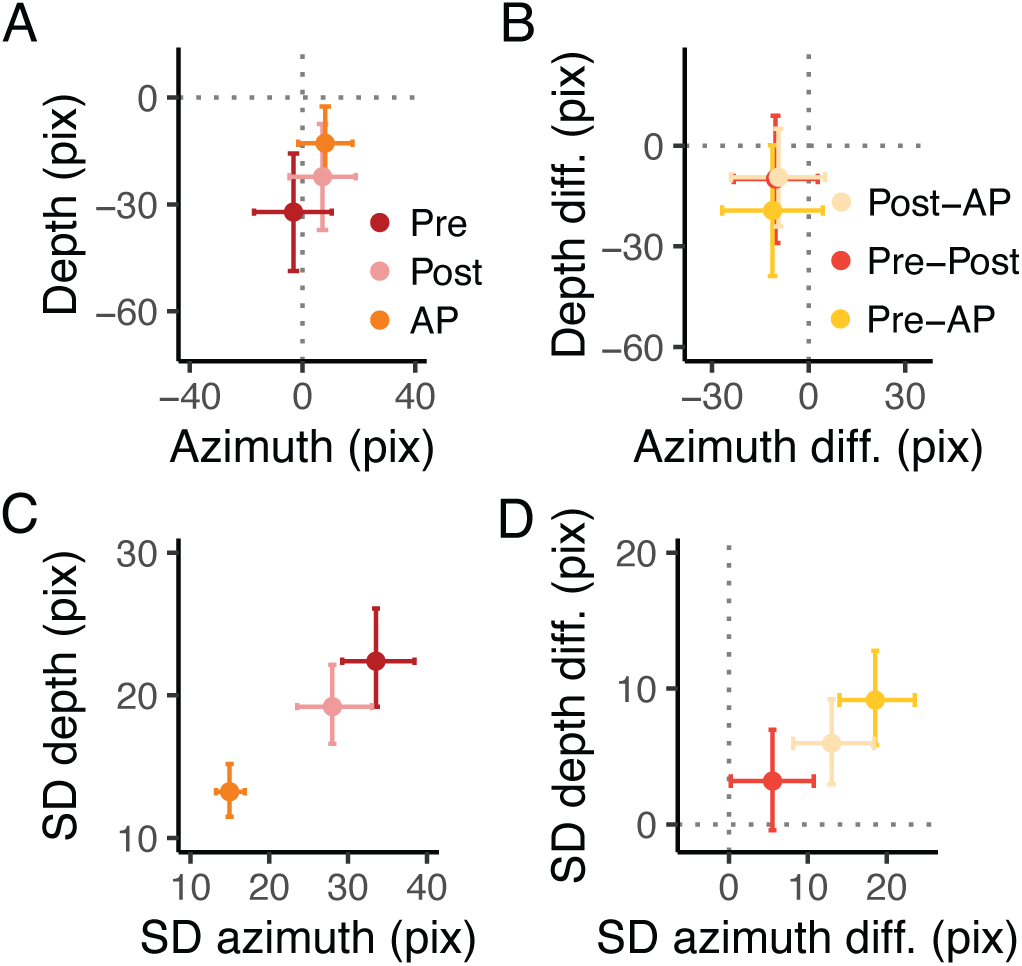
Experimental results. A) Estimates of the accuracy in the azimuth and depth. The dots represent the mean, and the error bars denote the 95% HDI of the estimates. The intersection of the grey dashed line represents the center of the targets. In A, positive values in the depth dimension indicate an overshoot, whereas positive values in the azimuth indicate that the index finger landed on the right side of the target. B) Estimates of the differences in accuracy in the azimuth and depth between conditions. The dots represent the mean, and the error bars denote the 95% HDI of the estimates. The grey dashed line represents the points of equal error in the depth and azimuth dimensions. C) Estimates of the SD in the azimuth and depth. The dots represent the mean, and the error bars denote the 95% HDI of the estimates. The intersection of the grey dashed line represents the center of the targets. D) Estimates of the differences in SD in the azimuth and depth between conditions. The dots represent the mean, and the error bars denote the 95% HDI of the estimates. The grey dashed line represents the points of equal variability in the depth and azimuth dimensions.

The SD instead (Figure 4B, Figure 5C and D), was credibly lower with a strong evidence toward the alternative hypothesis (BF_10_ *>*100) in Pre compared to AP (depth = 9.2, HDI = 5.84, 12.8 pixels; azimuth = 18.6, HDI = 14.0, 23.6 pixels) and in Post compared to AP (depth = 5.9, HDI = 2.9, 9.2 pixels; azimuth = 13.0, HDI = 8.1, 18.4 pixels). The SD between Pre and Post was credibly similar along the depth dimension (depth = 3.19, HDI = −0.4, 6.9 pixels), but not along the azimuth dimension (azimuth = 5.53., HDI = 0.2, 10.8). However, the BF showed very weak evidence toward the alternative hypothesis for all the dimensions (BF_10_ depth = 0.2, BF_10_ azimuth = 0.9). The slope for the SD was credibly lower than 0 in the depth dimension for the Pre (slope = −0.08, HDI = −0.1, −0.02 pixels) and Post conditions (slope = −0.07, HDI = −0.1,-0.02 pixels), with, however, anecdotal evidence toward the alternative hypothesis (BF_10_ *<*3). The comparison between conditions revealed a credibly lower slope in the depth dimension in the Pre compared to AP (slope diff. = −0.08, HDI = −0.1, −0.03 pixels) and in the Post compared to AP (slope diff. = −0.07, HDI = −0.1,-0.01 pixels). The Bayes factor analysis revealed, however, anecdotal evidence toward the alternative hypothesis (BF_10_ *<*3).

In sum, results showed that reaching an audio-proprioceptive target has a minimal impact on improving action performance toward auditory targets. Moreover, even though the SD in the Post was slightly lower than in the Pre condition, it did not reach the same level as in the AP condition.

#### Within Conditions Learning Effect

The analysis revealed a lack of learning effect (i.e., change in movement precision) across blocks (Figure 6A and B). The slope was not credibly different than 0 in each condition, revealing no changes in the SD as a function of the trial block. Additionally, comparison between conditions revealed a credible similar slope across conditions (Figure 6C). This further confirms that additional proprioceptive target information is insufficient to improve action performance throughout the task and that in the AP condition, participants mainly relied on proprioception to perform the movement.

**Figure 6:**
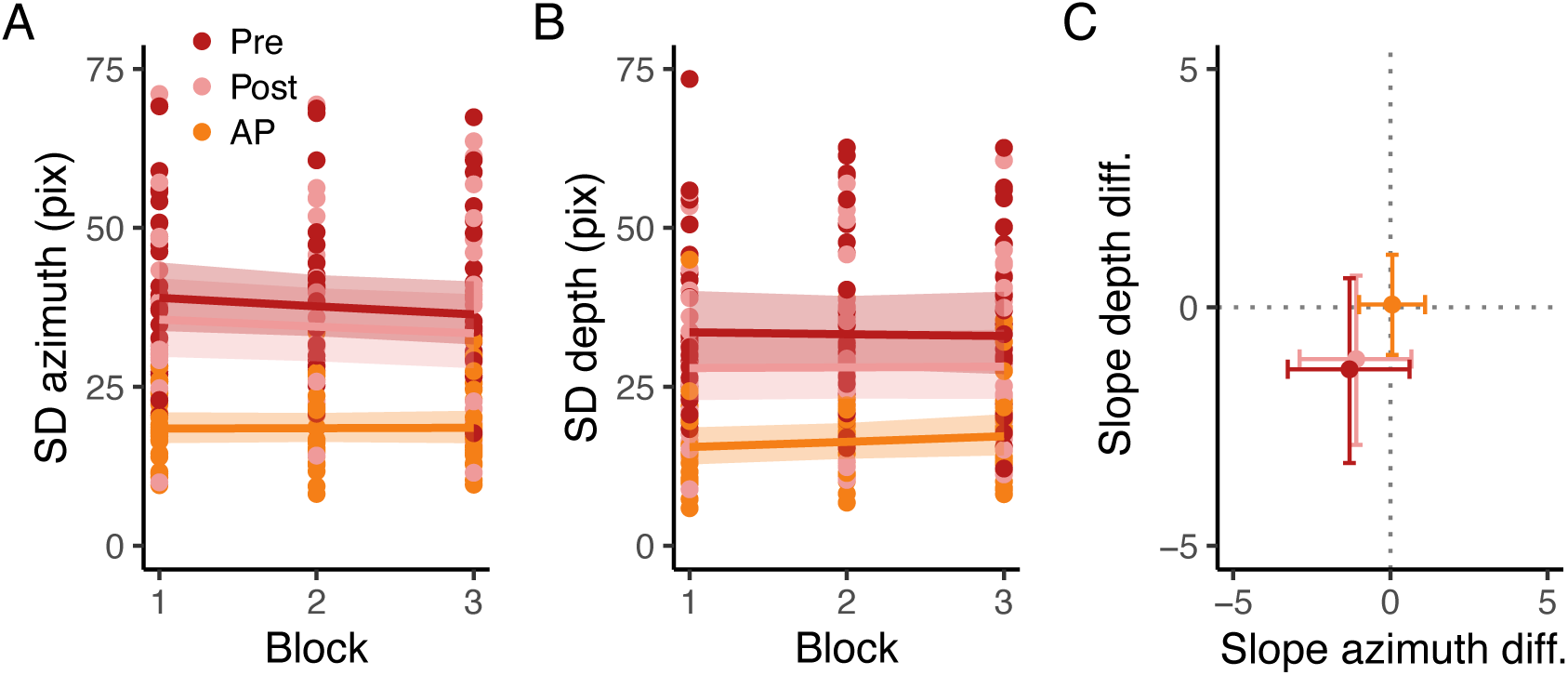
Learning effect results. A) Raw endpoint standard deviation in the azimuth and depth. The dots represent the raw single participant standard deviation, whereas solid lines and shaded regions represent the estimated standard deviation and the 95% credible intervals of the posterior distributions of the Bayesian regression model. B) Estimated slopes of the endpoint standard deviation as a function of the trial block for each condition. Points represent the mean and the error bars, the 95% credible intervals of the posterior distributions.

## Discussion

In this study, we investigated how reaching movements toward auditory and proprioceptive targets compare. To this aim, we asked participants to perform a reaching task toward an auditory, proprioceptive, or audio-proprioceptive target and measured the accuracy and precision (SD) in azimuth and depth dimensions. We found that actions toward proprioceptive and auditory targets share similarities in accuracy along the azimuth dimension, with slightly better accuracy for proprioception along the depth dimension. However, actions toward proprioceptive targets were more precise (lower SD) than those toward auditory targets in both azimuth and depth dimensions, thus disambiguating the results of previous studies that separately addressed auditory and proprioceptive target reaching performance (Parseihian et al, 2014; Maće et al, 2012; Boyer et al, 2013; Jones et al, 2012; Monaco et al, 2010; Khanafer and Cressman, 2014; Flannigan et al, 2018; McGuire and Sabes, 2009; Blouin et al, 2014; Cameron and Ĺopez-Moliner, 2015). Multisensory target reaching resembles proprioceptive target reaching for both accuracy and precision, confirming a high reliance on proprioception when reaching an audio-proprioceptive target (Lackner and Shenker, 1985).

Considering our results, the processes underpinning auditory and proprioceptive reaching may resemble those hypothesized by the Multimodal Theory of Sensory Integration for vision and haptics (Tagliabue and McIntyre, 2014). In the proprioceptive condition, both the *target* and the *reaching hand* positions were processed within the same sensory modality, allowing for a direct intra-modal comparison (Figure 7A). In contrast, in the auditory condition, the comparison involved different sensory modalities: the *target* location was specified via auditory input, while the position of the *reaching hand* remained available through proprioception (Figure 7B). This cross-modal comparison may have introduced additional variability in performance, leading to less consistent actions in the auditory condition compared to the proprioceptive one. In the audio-proprioceptive condition, proprioceptive signals conveyed information about both the *target* and the *reaching hand*, while auditory signals provided additional, redundant information about the *target* location. This involved an asymmetric distribution of sensory information, with proprioception contributing to both *target* and *reaching hand* localization, and audition contributing solely to the *target* (Figure 7C). Such asymmetry may have led to a predominant reliance on proprioceptive information to reduce action variability.

**Figure 7:**
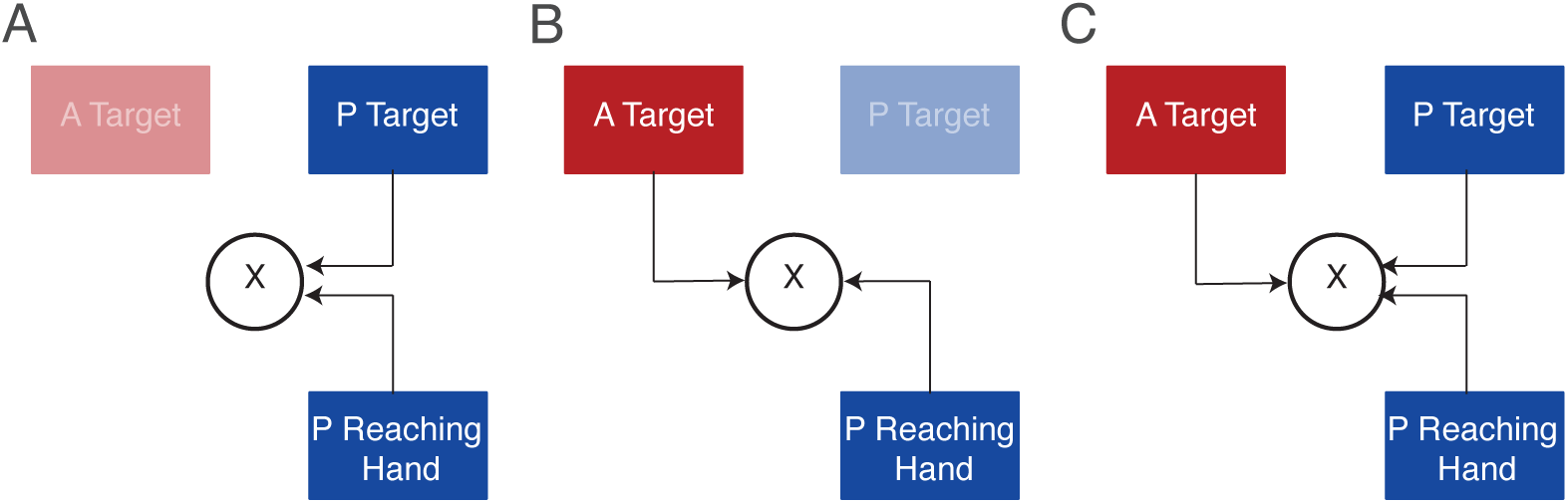
Schematic representation of the models of sensorimotor processes in unisensory and multisensory conditions. A) Combination of target and reaching hand information in the proprioceptive condition. B) Combination of target and reaching hand information in the auditory condition. C) Combination of auditory and proprioceptive target and proprioceptive reaching hand information in the audio-proprioceptive condition.

Although differing in precision, the comparable accuracy observed in both auditory and proprioceptive conditions in the azimuth dimension suggests the involvement of similar sensorimotor mechanisms across sensory modalities to accurately reach the target. Proprioceptive and tactile inputs from the reaching arm are inevitably engaged during interactions with sounds within reachable space (Serino, 2019; Farné and Làdavas, 2002; Van der Stoep et al, 2015). These interactions are underpinned by an integrated neural system that processes both sensory and motor information (Farné and Làdavas, 2002; Serino, 2019). Key components of this system include fronto-parietal areas such as the auditory cortex (Kayser et al, 2005; Caetano and Jousmäki, 2006; Zierul et al, 2017; Lohse et al, 2022), the ventral intraparietal area (Graziano et al, 1999; Graziano, 1999), and the premotor cortex (Serino, 2019), all of which are implicated in action planning and execution (Camponogara, 2023). Notably, activation within the motor cortex increases when sounds are presented within reachable space (Serino et al, 2009), enabling finely tuned motor responses (Canzoneri et al, 2012; Finisguerra et al, 2015; Noel et al, 2018; Serino, 2019; Camponogara et al, 2015; Bahadori et al, 2021; Bahadori and Cesari, 2021; Serino et al, 2009). Thus, reaching toward auditory targets may rely on specialized sensorimotor processes involving cortical networks processing inputs from both the *target* and the *reaching hand*, enabling comparable action accuracy to that observed in proprioceptive target reaching. However, such sensorimotor processes may not be sufficient to ensure consistent motor performance across movements in the auditory condition. The increased variability in the auditory condition may reflect the added complexity of the sensorimotor transformation required for localizing and acting upon auditory stimuli, where inputs from the auditory target are compared with proprioceptive information from the reaching hand, as previously discussed (Figure 7).

The potential benefit of proprioceptive information on auditory target reaching was investigated in a second experiment. Here, participants performed an auditory target reaching task both before and after an audio-proprioceptive target reaching session. While the effect was not strong, results revealed a slight reduction in variability along both movement dimensions. This modest improvement contrasts with previous studies reporting stronger effects of sensorimotor training on auditory localization. Several methodological differences between the current study and previous research may account for this discrepancy. Notably, many previous studies assessed participants’ auditory localization abilities under conditions of auditory disruption, either by including individuals with hearing impairments (Alzaher et al, 2023; Valzolgher et al, 2023b,a), simulating monaural hearing loss in normally hearing individuals (Valzolgher et al, 2020a,b, 2022, 2024), or manipulating the spectral content of auditory stimuli (Parise et al, 2024; Hofman, 1998; Isaiah et al, 2014). In such conditions, a training session consisting of reaching the auditory target has been shown to significantly enhance subsequent auditory localization performance (Isaiah et al, 2014; Parise et al, 2024; Valzolgher et al, 2022, 2024). In contrast, studies involving participants with intact hearing abilities typically report minimal or no improvements following auditory target reaching (Finocchietti et al, 2017; Cuppone et al, 2019). This suggests that training-related gains may emerge in disrupted auditory conditions when auditory reliability is compromised. The lack of substantial effects in the present study may reflect the relatively high baseline reliability of auditory inputs in our intact-hearing participants. Such effects may be more likely to emerge under conditions of impaired or degraded auditory perception.

An additional important consideration potentially explaining the small benefits of additional proprioceptive target information in our experiment relates to the use of terminal feedback in prior studies, which often involved visual (Fletcher and Zgheib, 2020; Fletcher et al, 2021a,b), audiovisual (Valzolgher et al, 2020a,b, 2023b,a, 2024), or proprioceptive and tactile cues (Hüg et al, 2019, 2022) to inform participants of their performance throughout the training phase. For example, in studies using virtual reality, participants were presented with an array of virtual speakers. During the pre- and post-training assessments, a sound was played, and participants were asked to verbally identify the corresponding speaker or to point their head toward the speaker emitting the sound. During the training phase, participants performed a reaching movement toward the sound-emitting speaker, which continued to play until the target was successfully reached, at which point the speaker turned green and the sound ceased (Valzolgher et al, 2020a,b, 2023b,a, 2024). Similarly, in studies involving physical speakers, blindfolded participants were instructed to reach and touch the speaker emitting the sound (Cuppone et al, 2018, 2019; Hüg et al, 2019, 2022). In this case, the tactile information participants experienced at the end of the movement acted as feedback about the final speaker position. It is well established that terminal visual (Hinder et al, 2008; Park et al, 2000; Guadagnoli and Kohl, 2001) and proprioceptive feedback (Müller et al, 2023) can drive sensorimotor adaptation (Taylor et al, 2014). The lack of any externally delivered terminal feedback in our study highlighted the specific role of additional proprioceptive target information in subsequent auditory reaching performance.

Taken together, our results show that action performance toward auditory and proprioceptive targets demonstrates comparable accuracy along the azimuth dimension, with proprioception showing slightly greater accuracy along the depth dimension. However, actions differ in precision, with those directed toward proprioceptive targets being more precise than those directed toward auditory targets. In multisensory target reaching, actions heavily rely on proprioceptive target information to reduce action uncertainty. When testing the effect of additional proprioceptive inputs on a subsequent auditory target reaching performance, the slight improvement in the auditory target reaching suggests that proprioceptive inputs could contribute to reducing the variability in the auditory target location, but such an effect is not as strong as when auditory information is disrupted (Alzaher et al, 2023; Valzolgher et al, 2023b,b, 2020a,b, 2022, 2024; Parise et al, 2024; Hofman, 1998; Isaiah et al, 2014).

### Limitations

There are several limitations to consider in the performed experiments, which are related to the use of a perforated board and the number of targets used. Even though perforated, the use of a wooden board may have attenuated the high-frequency spectrum, potentially reducing auditory reliability. Nevertheless, studies showed that precision in sound localization in the horizontal dimension decreased for limited-band high-frequency noises (3 – 10 kHz) compared to those within the low-frequency band (0.1 – 1 kHz) (Dobreva et al, 2011), thus suggesting a preserved precision in the auditory condition. The higher reliance on proprioception in experiment 1 and the slight improvement seen in experiment 2 could also be attributed to the limited number of target locations (three). It is possible that, in each trial, participants relied on the memory of the left-hand position to map the targets locations, leveraging this mapping to either enhance their performance in P and AP (experiment 1) or in the post-training session (experiment 2). An alternative explanation for the improvement in experiment 2 could involve the attenuated sound frequencies produced by the perforated wooden board, which may have reduced the reliability of auditory cues and enhanced reliance on proprioceptive cues. Even though studies showed that sound localization variability is lower (i.e., higher precision) for low compared to high frequencies sounds (Dobreva et al, 2011), a potential reduction in auditory reliability, combined with a limited number of auditory targets, may have led participants to increasingly depend on the previously sensed proprioceptive target positions to reach the auditory target in the post-training session.

## CRediT author statement

### Ivan Camponogara

Conceptualization, Methodology, Software, Validation, Formal analysis, Investigation, Data Curation, Writing - Original Draft, Writing - Review and Editing, Visualization, Supervision, Funding acquisition.

## Acknowledgements

We acknowledge the support of the Zayed University Research Incentive Fund (grant 23076).

